# Increased lung cell entry of B.1.617.2 and evasion of antibodies induced by infection and BNT162b2 vaccination

**DOI:** 10.1101/2021.06.23.449568

**Authors:** Prerna Arora, Amy Kempf, Inga Nehlmeier, Anzhalika Sidarovich, Nadine Krüger, Luise Graichen, Anna-Sophie Moldenhauer, Martin S. Winkler, Sebastian Schulz, Hans-Martin Jäck, Metodi V. Stankov, Georg M. N. Behrens, Stefan Pöhlmann, Markus Hoffmann

## Abstract

The delta variant of SARS-CoV-2, B.1.617.2, emerged in India and has subsequently spread to over 80 countries. B.1.617.2 rapidly replaced B.1.1.7 as the dominant virus in the United Kingdom, resulting in a steep increase in new infections, and a similar development is expected for other countries. Effective countermeasures require information on susceptibility of B.1.617.2 to control by antibodies elicited by vaccines and used for COVID-19 therapy. We show, using pseudotyping, that B.1.617.2 evades control by antibodies induced upon infection and BNT162b2 vaccination, although with lower efficiency as compared to B.1.351. Further, we found that B.1.617.2 is resistant against Bamlanivimab, a monoclonal antibody with emergency use authorization for COVID-19 therapy. Finally, we show increased Calu-3-lung cell entry and enhanced cell-to-cell fusion of B.1.617.2, which may contribute to augmented transmissibility and pathogenicity of this variant. These results identify B.1.617.2 as an immune evasion variant with increased capacity to enter and fuse lung cells.

## INTRODUCTION

Vaccines based on inactivated whole virus, adenoviral vectors or mRNAs encoding the severe acute respiratory syndrome coronavirus 2 (SARS-CoV-2) spike (S) protein protect against coronavirus disease 2019 (COVID-19) and allow to effectively combat the COVID-19 pandemic (Polack et al., 2020, Golob et al., 2021, Xia et al., 2021). These vaccines present the S proteins of viruses circulating early during the pandemic as antigens to the immune system. However, at a later stage of the pandemic so called SARS-CoV-2 variants of concern (VOC) emerged that harbor mutations in the S protein that allow for augmented transmissibility (B.1.1.7, alpha variant) and/or immune evasion (B.1.351, beta variant; P.1, gamma variant) (Plante et al., 2021b). Mutations conferring increased transmissibility might augment binding to the cellular receptor ACE2 while mutations conferring immune evasion alter epitopes of neutralizing antibodies (Plante et al., 2021b). Immune evasion can allow for infection of convalescent or vaccinated individuals but vectored and mRNA-based vaccines protect against severe COVID-19 induced by alpha, beta and gamma VOC.

A massive surge of COVID-19 cases was detected in India between April and May 2021 and was associated with spread of a new variant, B.1.617, that subsequently branched off into B.1.617.1, B.1.617.2 and B.1.617.3. The B.1.617.2 variant subsequently spread into more than 80 countries and became dominant in India and the United Kingdom(Singh et al., 2021, Campbell et al., 2021). In the UK, the spread of B.1.617.2 was associated with a marked increase in cases and more than 80% of new infections are now due to B.1.617.2. A rapid increase of B.1.617.2 spread is also expected in Germany, the US and several other countries, and a recent massive increase of cases in Lisbon, Portugal, that required travel restrictions is believed to be due to B.1.617.2. In order to contain spread of B.1.617.2, now considered a VOC, it will be critical to determine whether convalescent or vaccinated patients are protected against infection by this variant. Here, we addressed this question using reporter particles pseudotyped with the SARS-CoV-2 spike (S) protein, which are suitable tools to study SARS-CoV-2 neutralization by antibodies (Riepler et al., 2020, Schmidt et al., 2020).

## RESULTS

The S protein of B.1.617.2 harbors nine mutations in the surface unit, S1, of the S protein and 1 mutation in the transmembrane unit, S2 (Figure 1A-B). Mutations T19R, G142D, E156G, F157Δ and R158Δ are located in the N-terminal domain of S1, which contains epitopes for neutralizing antibodies (Liu et al., 2020, McCallum et al., 2021, Suryadevara et al., 2021, Chi et al., 2020). The receptor binding domain (RBD) harbors mutations L452R and T478K. Mutation L452R reduces antibody-mediated neutralization (Deng et al., 2021, Liu et al., 2021b) and it has been speculated that T478K might increase infectivity (Wang et al., 2021). Mutation D614G is located between RBD and S1/S2 cleavage sites and linked to increased ACE2 binding, replication in the upper respiratory tract and transmission (Figure 1A-B) (Plante et al., 2021a, Zhou et al., 2021). Finally, P681R might increase cleavage of S protein at the S1/S2 site while the impact of D950N on S protein driven entry and its inhibition by antibodies is unknown.

**Figure 1:**
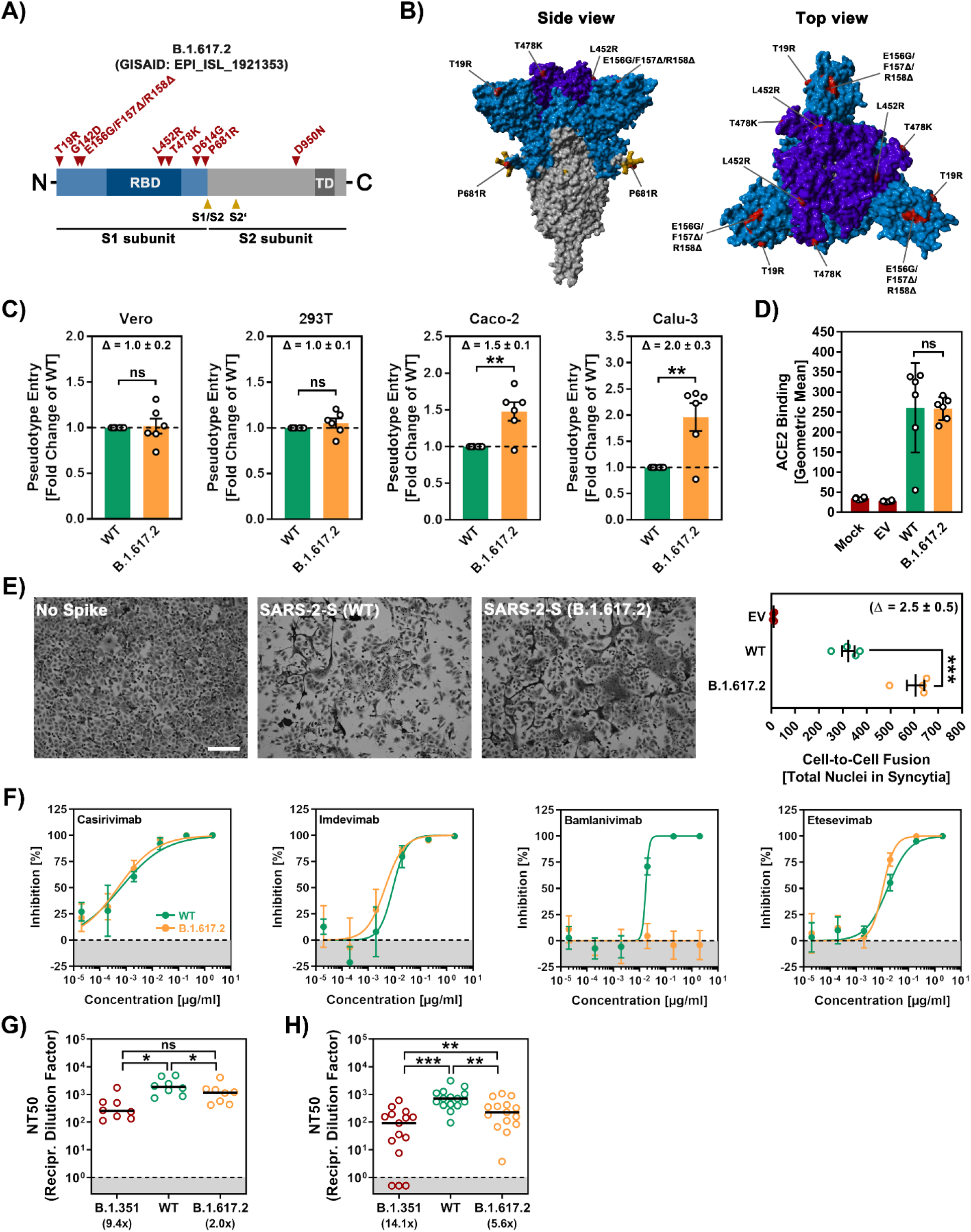
The spike protein of SARS-CoV-2 B.1.617.2 promotes efficient entry into human lung and colon cells, causes more cell-to-cell fusion and evades from antibody-mediated neutralization. (A) Schematic overview of the S protein from SARS-CoV-2 variant B.1.617.2 (RBD, receptor-binding domain; TD, transmembrane domain). (B) Location of the mutations found in SARS-CoV-2 variant B.1.617.2 in the context of the trimeric spike protein (Color code: light blue, S1 subunit with RBD in dark blue; gray, S2 subunit; orange, S1/S2 and S2’ cleavage sites; red, mutated amino acid residues). (C) Pseudotyped particles bearing the S protein of wildtype (WT) SARS-CoV-2 or variant B.1.617.2 were inoculated onto the indicated cell lines and transduction efficiency was quantified by measuring virus-encoded luciferase activity in cell lysates at 16-18 h post transduction. Presented are the average (mean) data from six biological replicates (each conducted with technical quadruplicates) for which transduction was normalized against SARS-CoV-2 S WT (= 1). Error bars indicate the standard error of the mean. Statistical significance of differences between WT and B.1.617.2 S proteins was analyzed by two-tailed Students t-test (p > 0.05, not significant [ns]; p ≤ 0.01, **). See also Figure S1A. (D) BHK-21 expressing the S protein of WT SARS-CoV-2 or variant B.1.617.2 were subsequently incubated with soluble ACE2 (harboring a C-terminal Fc-tag derived from human IgG) and AlexaFluor-488-conjugated anti-human antibody, before being subjected to flow cytometry. ACE2 binding efficiency was analyzed by measuring the geometric mean channel fluorescence at 488 nm. Untransfected cells and cells transfected with empty expression plasmid served as controls. Presented are the average (mean) data from six biological replicates (each conducted with single samples). Error bars indicate the standard deviation (SD). Statistical significance of differences between WT and variant B.1.617.2 S proteins was analyzed by two-tailed Students t-test (p > 0.05, ns). (E) Analysis of S protein induced cell-to-cell fusion. A549-ACE2 cells were transfected with expression plasmid for the indicated S proteins or empty vector (EV). At 24 h posttransfection, cells were fixed and subsequently stained with May-Gruenwald and Giemsa solutions. Presented are representative microscopic images (scale bar = 200 μm). For quantification of fusion efficiency, the total number of nuclei in syncytia per image was counted. Presented are the average (mean) data from four biological replicates (each conducted with single samples; for each sample, three randomly selected areas were imaged and independently analyzed by two persons). Error bars indicate the SEM. Statistical significance of differences between WT and B.1.617.2 S proteins was analyzed by two-tailed Students t-test (p ≤ 0.001, ***). (F) Neutralization of SARS-CoV-2 WT, B.1.351 and B.1.617.2 S proteins by monoclonal antibodies used for COVID-19 therapy. Pseudotyped particles bearing the S protein of WT SARS-CoV-2 or variant B.1.617.2 were incubated for 30 min at 37 °C in the presence of escalating concentrations (0.00002, 0.0002, 0.002, 0.02, 0.2, 2 μg/ml) of the indicated SARS-CoV-2 S protein-specific monoclonal antibody (please see Figure S1B) or an unrelated control antibody (please see Figure S1C), before being inoculated onto Vero cells. Transduction efficiency was quantified by measuring virus-encoded luciferase activity in cell lysates at 16-18 h post transduction. Presented are the average (mean) data from a single biological replicate (conducted with technical quadruplicates) for which transduction was normalized against samples that did not contain any antibody (= 0% inhibition). Error bars indicate the SD. (G) Neutralization of SARS-CoV-2 WT, B.1.351 and B.1.617.2 S proteins by antibodies in convalescent plasma. Pseudotyped particles bearing the S protein of WT SARS-CoV-2 or variant B.1.617.2 were incubated for 30 min at 37 °C in the presence of different dilutions of convalescent plasma (1:25, 1:100, 1:400, 1:1,600, 1:6,400, 1:25,600). Transduction efficiency was quantified by measuring virus-encoded luciferase activity in cell lysates at 16-18 h post-transduction and used to calculate the plasma dilution factor that leads to 50 % reduction in S protein-driven cell entry (neutralizing titer 50, NT50). Presented are the data from a total of eight convalescent plasma (black lines indicate the median). Statistical significance of differences between the indicated groups was analyzed by two-tailed Students t-test (p > 0.05, ns; p ≤ 0.05, *; p ≤ 0.01, **; p ≤ 0.001, ***). Please see also Figure S1D. (G) The experiment was performed as described for panel F but this time serum from Comirnaty/BNT162b2-vaccinated individuals was investigated. Presented are the data from a total of fifteen vaccinee sera (black lines indicate the median). Please see also Figure S1E.

We first asked whether B.1.617.2 S protein mediates robust entry into cell lines frequently used for SARS-CoV-2 research, Vero (African green monkey, kidney), 293T (human, kidney), Caco-2 (human, colon) and Calu-3 (human, lung). All cell lines express endogenous ACE2 and Vero, Caco-2 and Calu-3 cells are often used for infection studies with authentic SARS-CoV-2. The B.1.617.2 S protein mediated entry into 293T and Vero cells with the same efficiency as WT S protein while entry into Caco-2 (~1.5-fold) and Calu-3 cells (~2.0-fold) was augmented (Figure 1C and Supplemental figure 1A). The lung is the central target of SARS-CoV-2 but infection of colon has also been reported, suggesting that B.1617.2 might have increased capacity to enter target cells in these tissues. Finally, we did not detect increased ACE2 binding of B.1.617.2 S protein (Figure 1D), suggesting that increased entry into Caco-2 and Calu-3 cells was not due to augmented ACE2 binding.

Besides its ability to drive fusion of viral and cellular membranes, the S protein is further able to drive the fusion of neighboring cells, resulting in the formation of multinucleated giant cells, so called syncytia, which have been observed in vitro following directed S protein expression or SARS-CoV-2 infection and in post mortem tissues from COVID-19 patients (Bussani et al., 2020, Tian et al., 2020, Xu et al., 2020) . Since SARS-CoV-2 S protein-driven syncytium formation is believed to contribute to COVID-19 pathogenesis, we investigated the ability of B.1.617.2 S protein to drive cell-to-cell fusion in the human lung cell line A549, which was engineered to express high levels of ACE2. As expected, directed expression of WT S led to the formation of syncytia, while cells transfected with empty expression plasmid remained normal (Figure 1E). Strikingly, directed expression of B.1.617.2 S protein caused more and larger syncytia and quantification of cell-to-cell fusion revealed that fusion by B.1.617.2 S protein was ~2.5-fold more efficient as compared to WT S (Figure 1E).

We next determined whether entry of B.1.617.2 is susceptible to inhibition by recombinant antibodies with emergency use authorization for COVID-19 treatment. Three out four antibodies tested inhibited B.1.617.2 S protein with similar efficiency as WT S protein (Figure 1F and Supplemental figure 1C). However, B.1.617.2 was resistant to Bamlanivimab, most likely because of mutation L452R (Supplemental figure 1B and (Starr et al., 2021)). Thus, Bamlanivimab monotherapy is not suitable for prevention or treatment of B.1.617.2 infection. Finally, we asked whether B.1.617.2 entry is inhibited by antibodies generated by infected or vaccinated individuals. For these experiments, we employed the S protein of B.1.351 as control since this VOC exhibits marked evasion from neutralizing antibodies. A previously described collection of plasma (Hoffmann et al., 2021a) from convalescent COVID-19 patients collected at University Hospital Göttingen, Germany, neutralized entry driven by B.1.617.2 S protein with slightly reduced efficiency as compared to WT S protein (Figure 1G and Supplemental figure 1D). In contrast, neutralization of B.1.351 S protein-dependent entry was markedly reduced. Finally, similar observations were made with previously characterized sera (Hoffmann et al., 2021b) from donors who received two doses of BNT162b2, although immune evasion of B.1.617.2. was more prominent as compared to convalescent sera (Figure 1H and Supplemental figure 1E).

## DISCUSSION

Our results demonstrate immune evasion, enhanced colon- and lung cell entry and augmented syncytium formation by B.1.617.2. Evasion of antibody-mediated neutralization by B.1.617.2 is in agreement with two recent studies (Liu et al., 2021a, Wall et al., 2021) and is more prominent than previously observed by us for B.1.1.7 but less prominent as compared to B.1.351 (Hoffmann et al., 2021a). This finding would be compatible with increased vaccine breakthrough of B.1.617.2 but also suggests that BNT162b2 should still protect from B.1.617.2-induced COVID-19. Treatment of infection with Bamlanivimab alone will be ineffective but we expect that Casirivimab, Imdevimab and Etesevimab will be beneficial to B.1.617.2 infected patients when administered early after infection. The observation that B.1.617.2 S protein is able to cause more cell-to-cell fusion than WT S may suggest that B.1.617.2 could cause more tissue damage, and thus be more pathogenic, than previous variants or that viral spread via syncytium formation contributes to efficient inter- and intra-host spread of this variant. Entry experiments with cell lines need to be interpreted with care and confirmation with primary cells is pending. However, the significantly increased entry into the colon and lung cell lines Caco-2 and Calu-3, respectively, suggest that B.1.617.2 might have an augmented capacity to infect these organs and increased infection of the respiratory epithelium might account for the purported increased transmissibility of B.1.617.2.

## AUTHOR CONTRIBUTIONS

Conceptualization, M.H., S.P.; Funding acquisition, S.P.; Investigation, P.A., A.K., I.N., A.S., N.K., L.G., A.-S.M., M.H.; Essential resources, M.S.W., S.S., H.-M.J., M.V.S., G.M.N.B.; Writing, M.H., S.P.; Review and editing, all authors.

## ACKNOWLEDGMENTS

We like to thank Roberto Cattaneo, Georg Herrler, Stephan Ludwig, Andrea Maisner, Thomas Pietschmann and Gert Zimmer for providing reagents. We gratefully acknowledge the originating laboratories responsible for obtaining the specimens and the submitting laboratories where genetic sequence data were generated and shared via the GISAID Initiative, on which this research is based. S.P. acknowledges funding by BMBF (01KI2006D, 01KI20328A, 01KI20396, DM11-311), the Ministry for Science and Culture of Lower Saxony (DM11-607, DM11-610, DM11-611) and the German Research Foundation (DFG; PO 716/11-1, PO 716/14-1). N.K. acknowledges funding by BMBF (ANI-CoV, 01KI2074A). M.S.W. received unrestricted funding from Sartorius AG, Lung research.

## COMPETING INTERESTS

The authors declare no competing interests

## MATERIALS AND METHODS

### Cell culture

All cell lines were incubated at 37 °C in a humidified atmosphere containing 5% CO_2_. 293T (human, female, kidney; ACC-635, DSMZ, RRID: CVCL_0063), Vero76 cells (African green monkey kidney, female, kidney; CRL-1586, ATCC; RRID: CVCL_0574, kindly provided by Andrea Maisner) and BHK-21 (Syrian hamster, male, kidney; ATCC Cat# CCL-10; RRID: CVCL_1915, kindly provided by Georg Herrler) and A549-ACE2 cells (Hoffmann et al., 2021a), which were derived from parental A549 cells (human, male, lung; CRM-CCL-185, ATCC, RRID:CVCL_0023; kindly provided by Georg Herrler), were cultured in Dulbecco’s modified Eagle medium (PAN-Biotech) supplemented with 10% fetal bovine serum (FBS, Biochrom), 100 U/ml penicillin and 0.1 mg/ml streptomycin (pen/strep) (PAN-Biotech). In addition, Calu-3 (human, male, lung; HTB-55, ATCC; RRID: CVCL_0609, kindly provided by Stephan Ludwig) and Caco-2 cells (human, male, colon; HTB-37, ATCC, RRID: CVCL_0025) were cultured in minimum essential medium (GIBCO) supplemented with 10% FBS, 1% pen/strep, 1x non-essential amino acid solution (from 100x stock, PAA) and 1 mM sodium pyruvate (Thermo Fisher Scientific). Cell lines were validated by STR-typing, amplification and sequencing of a fragment of the cytochrome c oxidase gene, microscopic examination and/or according to their growth characteristics. Furthermore, all cell lines were routinely tested for contamination by mycoplasma contamination.

### Expression plasmids

Plasmids encoding pCAGGS-DsRed, pCAGGS-VSV-G (vesicular stomatitis virus glycoprotein), pCG1-WT SARS-CoV-2 S (codon optimized, based on the Wuhan/Hu-1/2019 isolate, equipped with D614G mutation; with C-terminal truncation of the last 18 amino acid), pCG1-SARS-CoV-2 S, B.1.351 (codon optimized; with C-terminal truncation of the last 18 amino acid), ACE2 (angiotensin converting enzyme 2) and soluble ACE2 have been previously described (Hoffmann et al., 2021a, Hoffmann et al., 2021b, Hoffmann et al., 2020). In order to generate the expression vector for the S protein of SARS-CoV-2 variant B.1.617.2, the respective mutations were inserted into the WT SARS-CoV-2 S sequence by splice-overlap PCR. The resulting open reading frame was further inserted into vector pCG1 plasmid (kindly provided by Roberto Cattaneo, Mayo Clinic College of Medicine, Rochester, MN, USA), using BamHI and XbaI restriction enzymes. The integrity of all sequences was confirmed by sequence analysis using a commercial sequencing service (Microsynth SeqLab). Specific details on the cloning procedure can be obtained upon request. Transfection of 293T cells was carried out by the calcium-phosphate precipitation method, while BHK-21 and A549-ACE2 cells were transfected using Lipofectamine LTX (Thermo Fisher Scientific).

### Sequence analysis and protein models

The S protein sequence of SARS-CoV-2 S variant B.1.617.2 (GISAID Accession ID: EPI_ISL_1921353) was obtained from the GISAID (global initiative on sharing all influenza data) databank (https://www.gisaid.org/). Protein models were generated employing the YASARA software (http://www.yasara.org/index.html) and are based on published crystal structure PDB: 6XDG (Hansen et al., 2020), PDB: 7L3N (Jones et al., 2020) or PDB: 7C01 (Shi et al., 2020), or a template that was constructed by modelling the SARS-2 S sequence on PDB: 6XR8 (Cai et al., 2020), using the SWISS-MODEL online tool (https://swissmodel.expasy.org)

### Production of pseudotype particles

Rhabdoviral pseudotypes bearing SARS-CoV-2 spike protein were generated according to an established protocol (Berger Rentsch and Zimmer, 2011). Briefly, 293T cells were transfected with expression plasmids encoding S protein, VSV-G or empty plasmid (control). At 24 h posttransfection, cells were inoculated with a replication-deficient vesicular stomatitis virus that lacks the genetic information for VSV-G and instead codes for two reporter proteins, enhanced green fluorescent protein and firefly luciferase (FLuc), VSV∗ΔG-FLuc (kindly provided by Gert Zimmer) at a multiplicity of infection of 3. Following 1 h of incubation at 37 °C, the inoculum was removed and cells were washed with phosphate-buffered saline (PBS). Subsequently, cells received culture medium containing anti-VSV-G antibody (culture supernatant from I1-hybridoma cells; ATCC no. CRL-2700; except for cells expressing VSV-G, which received only medium) in order to neutralize residual input virus. After 16-18h, the culture supernatant was harvested, clarified from cellular debris by centrifugation at 4,000 x g, 10 min, aliquoted and stored at −80 °C.

### Transduction of target cells

For transduction experiments, target cells were seeded in 96-well plates and inoculated with equal volumes of pseudotype particles. The transduction efficiency was evaluated at 16-18 h post transduction. For this, cells were lysed in PBS containing 0.5% triton X-100 (Carl Roth) for 30 min at RT. Afterwards, cell lysates were transferred into white 96-well plates and mixed with luciferase substrate (Beetle-Juice, PJK) before luminescence was recorded using a Hidex Sense Plate luminometer (Hidex).

### Analysis of ACE2 binding

For the production of soluble ACE2 fused to the Fc portion of human immunoglobulin G (IgG), sol-ACE2, 293T cells were seeded in a T-75 flask and transfected with 20 μg of sol-ACE2 expression plasmid. The medium was replaced at 10 h posttransfection and cells were further incubated for 38 h. Further, the culture supernatant was harvested and clarified by centrifugation at 2,000 x g, 10 min, 4 °C. Next, the clarified supernatant was loaded onto Vivaspin protein concentrator columns (molecular weight cut-off of 30 kDa; Sartorius) and centrifuged at 4,000 x g at 4 °C until the supernatant was 100-fold concentrated. Finally, concentrated sol-ACE2 was aliquoted and stored at −80 °C.

In order to test the binding efficiency of sol-ACE2 to S protein, BHK-21 cells were seeded in 12-well plates and transfected with expression plasmid for WT or SARS-CoV-2 S variant. Untransfected cells and cells transfected with empty pCG1 plasmid served as controls. At 24 h posttransfection, the culture supernatant was removed and cells were washed and resuspended in PBS and transferred into 1.5 ml reaction before being pelleted by centrifugation (600 x g, 5 min, RT, all centrifugation steps). Thereafter, cells were washed with PBS containing 1% bovine serum albumin (BSA; PBS/BSA) and pelleted again by centrifugation. Next, the supernatant was removed and cell pellets were incubated with 100 μl of solACE2-Fc (1:100 in PBS/BSA) and rotated for 1 h at 4 °C using a Rotospin eppi rotator disk (IKA). After incubation, cells were pelleted and incubated with 100 μl of human AlexaFlour-488-conjugated antibody (1:200 in PBS/BSA; Thermo Fisher Scientific) and rotated again as described above. Finally, cells were washed and resuspended in PBS/BSA and subjected to flow cytometry using a LSR II flow cytometer and the FACS diva software (BD Biosciences). Data analysis was performed using the FCS express 4 Flow research software (De Novo Software) in order to obtain the geometric mean values.

### Syncytium formation assay

In order to analyze S protein-driven cell-to-cell fusion, A549-ACE2 cells were grown in 12-well plates and transfected with expression vector for WT or B.1.617.2 S protein. In addition, cells transfected with empty plasmid served as control. At 24 h posttransfection, cells were washed with PBS and fixed by incubation (20 min, room temperature) with 4% paraformaldehyde solution (Carl Roth). Thereafter, cells were washed with deionized water, air-dried and stained with May-Gruenwald solution (30 min, room temperature; Sigma-Aldrich). Next, cells were washed three times with deionized water, air-dried and 1:10 diluted Giemsa (30 min, room temperature; Sigma-Aldrich) solutions. Finally, cells were washed three times with deionized water, air-dried and analyzed by bright-field microscopy using a Zeiss LSM800 confocal laser scanning microscope and the ZEN imaging software (Zeiss). For each sample, three randomly selected areas were imaged and S protein-driven syncytium formation was quantified by counting the total number of nuclei in syncytia per image. Syncytia were defined as cells containing at least three nuclei. To eliminate potential bias and correct for counting errors, counting was performed blinded by two persons independently and for each sample average counts were used. Further, for each biological replicate, the average (mean) total number of nuclei in syncytia per image was calculated from three images obtained for randomly selected areas of the well.

### Collection of serum and plasma samples

Before analysis, all serum and plasma samples were heat-inactivated at 56 °C for 30 min. Further, all plasma/serum samples were pre-screened for their ability to neutralize transduction of Vero cells by pseudotype particles bearing WT SARS-CoV-2 S.

Convalescent plasma was obtained from COVID-19 patients treated at the intensive care unit of the University Medicine Göttingen (UMG) under approval given by the ethic committee of the UMG (SeptImmun Study 25/4/19 Ü). Cell Preparation Tube (CPT) vacutainers with sodium citrate were used for collection of convalescent plasma. Further, plasma was collected as supernatant over the peripheral blood mononuclear cell layer. In addition to convalescent plasma, serum from individuals vaccinated with BioNTech/Pfizer vaccine BNT162b2/Comirnaty was collected 24-31 days after receiving the second dose using S-Monovette® EDTA tubes (Sarstedt). Sampling and sample analysis were approved by the Institutional Review Board of Hannover Medical School (8973_BO_K_2020, amendment Dec 2020).

### Neutralization assay

For neutralization experiments, S protein bearing pseudotype particles were pre-incubated for 30 min at 37 °C with different concentrations of Casirivimab, Imdevimab, Bamlanivimab, Etesevimab, or unrelated control IgG (2, 0.2, 0.02, 0.002, 0.0002, 0.00002 μg/ml). Alternatively, pseudotype particles were pre-incubated with different dilutions (1:25, 1:100, 1:400, 1:1,600, 1:6,400) of convalescent plasma or serum from BNT162b2/Comirnaty vaccinated individuals. Following incubation, mixtures were inoculated onto Vero cells with particles incubated only with medium serving as control. Transduction efficiency was determined at 16-18 h postinoculation as described above.

### Data analysis

The results on S protein-driven cell entry represent average (mean) data acquired from six biological replicates, each conducted with four technical replicates. Data were normalized against WT S protein, for which entry was set as 1. Alternatively, transduction was normalized against the background signal (luminescence measured for cells inoculated with particles bearing no viral glycoprotein; set as 1). SolACE2 binding results are average (mean) data obtained from six biological replicates, each conducted with single samples. Each data point represents the geometric mean channel fluorescence for one biological replicate without normalization. Data on S protein-driven cell-to-cell fusion represent average (mean) data from four biological replicates, each conducted with single samples (three images per sample). Each data point represents the average (mean) number of nuclei in syncytia per image from two counting events by independent persons for each of the four biological replicate without normalization. The neutralization data are based on a single experiment (standard in the field), which were conducted with technical quadruplicates. For data normalized, background signals (Fluc signals obtained from cell inoculated with pseudotype particles bearing no S protein) were subtracted from all values and transduction by particles incubated only with medium was set as 0% inhibition. The neutralizing titer 50 (NT50) value, which indicates the plasma/serum dilution that causes a 50 % reduction of transduction efficiency, was calculated using a non-linear regression model (inhibitor vs. normalized response, variable slope).

Error bars are defined as either standard deviation (SD, neutralization data) or standard error of the mean (SEM, all other data). Data ware analyzed using Microsoft Excel (as part of the Microsoft Office software package, version 2019, Microsoft Corporation) and GraphPad Prism 8 version 8.4.3 (GraphPad Software). Statistical significance was tested by two-tailed Students t-test. Only p values of 0.05 or lower were considered statistically significant (p > 0.05, not significant [ns]; p ≤ 0.05, *; p ≤ 0.01, **; p ≤ 0.001, ***). Details on the statistical test and the error bars can be found in the figure legends.

## SI Figure Legend

**Figure S1:**
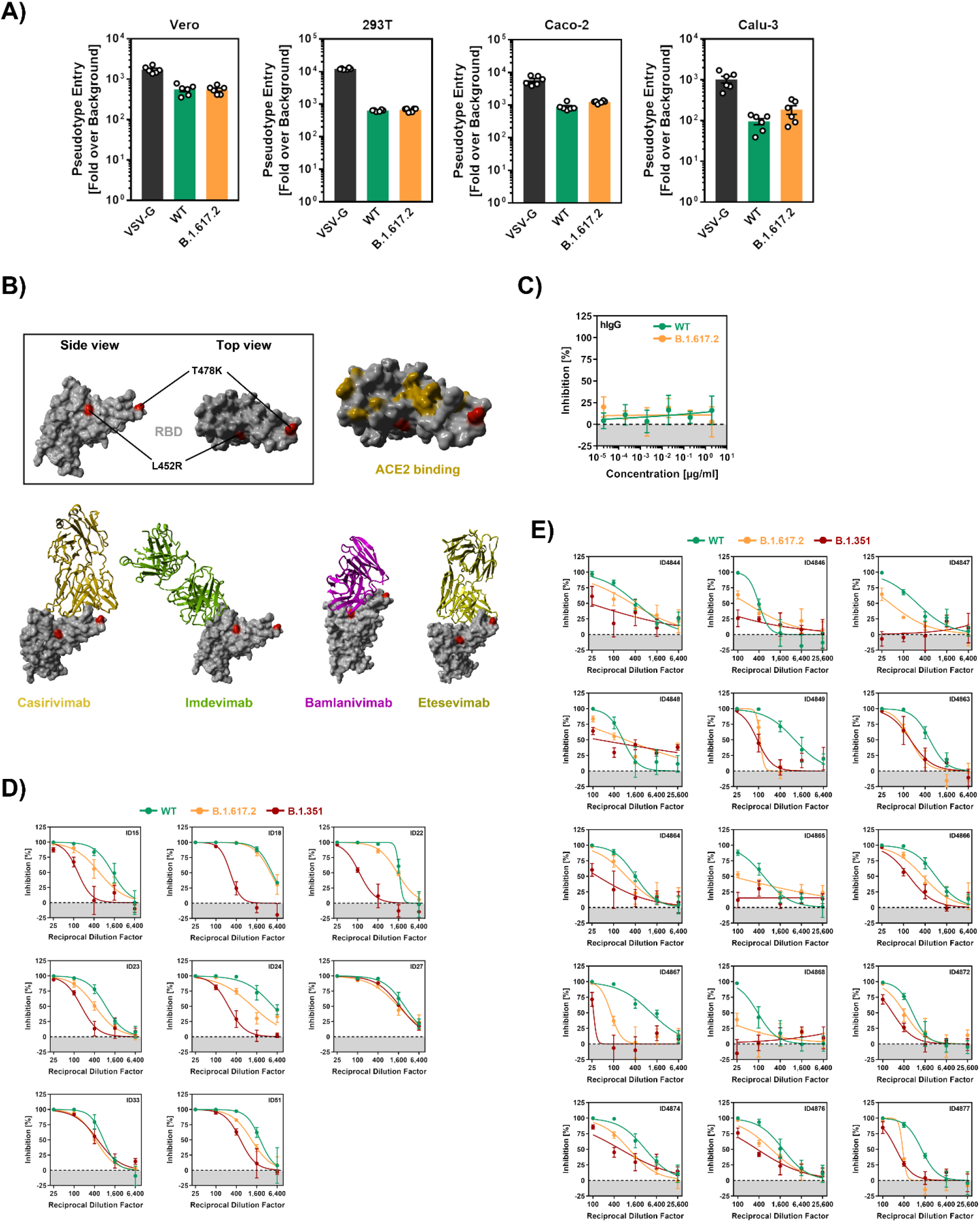
Cell entry and evasion of antibody-mediated neutralization by the spike protein of SARS-CoV-2 B.1.617.2 (related to Figure 1). (A) Transduction data normalized against the assay background (related to Figure 1C). The experiment was performed as described in the legend of Figure 1C. Presented are the average (mean) data from the same six biological replicates (each conducted with technical quadruplicates) as presented in Figure 1C with the difference that transduction was normalized against signals obtained from cells inoculated with particles bearing no viral glycoprotein (background, set as 1). In addition, transduction data of particles bearing VSV-G are included. Error bars indicate the SEM. (B) Location of the receptor binding domain (gray) mutations L452R and T478K (both red) of SARS-CoV-2 variant B.1.617.2 in the context of the interfaces for ACE2 binding (orange) and binding of monoclonal antibodies used for COVID-19 therapy. (C) An unrelated control antibody does not affect cell entry of pseudotype particles bearing SARS-CoV-2 WT, B.1.351 or B.1.617.2 S (related to Figure 1E). The experiment was performed as described in the legend of Figure 1E. (D) Individual neutralization data for convalescent plasma (related to Figure 1F). Pseudotype particles bearing the indicated S proteins were incubated (30 min, 37 °C) with different dilutions of convalescent plasma before being inoculated onto Vero cells. Transduction efficiency was quantified by measuring virus-encoded luciferase activity in cell lysates at 16-18 h posttransduction. Presented are the data from a single representative experiment conducted with technical quadruplicates. For normalization, inhibition of S protein-driven entry in samples without plasma was set as 0%. Error bars indicate the SD. The data were further used to calculated the plasma/serum dilution that leads to 50% reduction in S protein-driven cell entry (neutralizing titer 50, NT50; shown in Figure 1F). (E) Individual neutralization data for vaccinee serum (related to Figure 1G). Pseudotype particles bearing the indicated S proteins were incubated (30 min, 37 °C) with different dilutions of serum from individuals vaccinated with the Pfizer/BioNTech vaccine Comirnaty/BNT162b2 before being inoculated onto Vero cells. Transduction efficiency was quantified by measuring virus-encoded luciferase activity in cell lysates at 16-18 h posttransduction. Presented are the data from a single representative experiment conducted with technical quadruplicates. For normalization, inhibition of S protein-driven entry in samples without plasma was set as 0%. Error bars indicate the SD. The data were further used to calculated the NT50 shown (shown in Figure 1G).

## Notes

### Competing Interest Statement

The authors have declared no competing interest.

## REFERENCES

Berger Rentsc, M. & Zimmer, G. 2011. A vesicular stomatitis virus replicon-based bioassay for the rapid and sensitive determination of multi-species type I interferon. PLoS One, 6, e25858.

Bussani, R., Schneider, E., Zentilin, L., Collesi, C., Ali, H., Braga, L., Volpe, M. C., Colliva, A., Zanconati, F., Berlot, G., Silvestri, F., Zacchigna, S. & Giacca, M. 2020. Persistence of viral RNA, pneumocyte syncytia and thrombosis are hallmarks of advanced COVID-19 pathology. EBioMedicine, 61, 103104.

Cai, Y., Zhang, J., Xiao, T., Peng, H., Sterling, S. M., Walsh, R. M., JR., Rawson, S., Rits-Volloch, S. & Chen, B. 2020. Distinct conformational states of SARS-CoV-2 spike protein. Science, 369, 1586–1592.

Campbell, F., Archer, B., Laurenson-Schafer, H., Jinnai, Y., Konings, F., Batra, N., Pavlin, B., Vandemaele, K., Van Kerkhove, M. D., Jombart, T., Morgan, O. & Le Polain De Waroux, O. 2021. Increased transmissibility and global spread of SARS-CoV-2 variants of concern as at June 2021. Euro Surveill, 26.

Chi, X., Yan, R., Zhang, J., Zhang, G., Zhang, Y., Hao, M., Zhang, Z., Fan, P., Dong, Y., Yang, Y., Chen, Z., Guo, Y., Zhang, J., Li, Y., Song, X., Chen, Y., Xia, L., Fu, L., Hou, L., Xu, J., Yu, C., Li, J., Zhou, Q. & Chen, W. 2020. A neutralizing human antibody binds to the N-terminal domain of the Spike protein of SARS-CoV-2. Science, 369, 650–655.

Deng, X., Garcia-Knight, M. A., Khalid, M. M., Servellita, V., Wang, C., Morris, M. K., Sotomayor-Gonzalez, A., Glasner, D. R., Reyes, K. R., Gliwa, A. S., Reddy, N. P., Sanchez San Martin, C., Federman, S., Cheng, J., Balcerek, J., Taylor, J., Streithorst, J. A., Miller, S., Sreekumar, B., Chen, P. Y., Schulze-Gahmen, U., Taha, T. Y., Hayashi, J. M., Simoneau, C. R., Kumar, G. R., Mcmahon, S., Lidsky, P. V., Xiao, Y., Hemarajata, P., Green, N. M., Espinosa, A., Kath, C., Haw, M., Bell, J., Hacker, J. K., Hanson, C., Wadford, D. A., Anaya, C., Ferguson, D., Frankino, P. A., Shivram, H., Lareau, L. F., Wyman, S. K., Ott, M., Andino, R. & Chiu, C. Y. 2021. Transmission, infectivity, and neutralization of a spike L452R SARS-CoV-2 variant. Cell.

Golob, J. L., Lugogo, N., Lauring, A. S. & Lok, A. S. 2021. SARS-CoV-2 vaccines: a triumph of science and collaboration. JCI Insight, 6.

Hansen, J., Baum, A., Pascal, K. E., Russo, V., Giordano, S., Wloga, E., Fulton, B. O., Yan, Y., Koon, K., Patel, K., Chung, K. M., Hermann, A., Ullman, E., Cruz, J., Rafique, A., Huang, T., Fairhurst, J., Libertiny, C., Malbec, M., Lee, W. Y., Welsh, R., Farr, G., Pennington, S., Deshpande, D., Cheng, J., Watty, A., Bouffard, P., Babb, R., Levenkova, N., Chen, C., Zhang, B., Romero Hernandez, A., Saotome, K., Zhou, Y., Franklin, M., Sivapalasingam, S., Lye, D. C., Weston, S., Logue, J., Haupt, R., Frieman, M., Chen, G., Olson, W., Murphy, A. J., Stahl, N., Yancopoulos, G. D. & Kyratsous, C. A. 2020. Studies in humanized mice and convalescent humans yield a SARS-CoV-2 antibody cocktail. Science, 369, 1010–1014.

Hoffmann, M., Arora, P., Gross, R., Seidel, A., Hornich, B. F., Hahn, A. S., Kruger, N., Graichen, L., Hofmann-Winkler, H., Kempf, A., Winkler, M. S., Schulz, S., Jack, H. M., Jahrsdorfer, B., Schrezenmeier, H., Muller, M., Kleger, A., Munch, J. & Pohlmann, S. 2021a. SARS-CoV-2 variants B.1.351 and P.1 escape from neutralizing antibodies. Cell.

Hoffmann, M., Hofmann-Winkler, H., KrÜGer, N., Kempf, A., Nehlmeier, I., Graichen, L., Sidarovich, A., Moldenhauer, A.-S., Winkler, M. S., Schulz, S., JÄck, H.-M., Stankov, M. V., Behrens, G. M. N. & PÖHlmann, S. 2021b. SARS-CoV-2 variant B.1.617 is resistant to Bamlanivimab and evades antibodies induced by infection and vaccination. bioRxiv, 2021.05.04.442663.

Hoffmann, M., Kleine-Weber, H., Schroeder, S., Kruger, N., Herrler, T., Erichsen, S., Schiergens, T. S., Herrler, G., Wu, N. H., Nitsche, A., Muller, M. A., Drosten, C. & Pohlmann, S. 2020. SARS-CoV-2 Cell Entry Depends on ACE2 and TMPRSS2 and Is Blocked by a Clinically Proven Protease Inhibitor. Cell, 181, 271–280 e8.

Jones, B. E., Brown-Augsburger, P. L., Corbett, K. S., Westendorf, K., Davies, J., Cujec, T. P., Wiethoff, C. M., Blackbourne, J. L., Heinz, B. A., Foster, D., Higgs, R. E., Balasubramaniam, D., Wang, L., Bidshahri, R., Kraft, L., Hwang, Y., Zentelis, S., Jepson, K. R., Goya, R., Smith, M. A., Collins, D. W., Hinshaw, S. J., Tycho, S. A., Pellacani, D., Xiang, P., Muthuraman, K., Sobhanifar, S., Piper, M. H., Triana, F. J., Hendle, J., Pustilnik, A., Adams, A. C., Berens, S. J., Baric, R. S., Martinez, D. R., Cross, R. W., Geisbert, T. W., Borisevich, V., Abiona, O., Belli, H. M., De Vries, M., Mohamed, A., Dittmann, M., Samanovic, M., Mulligan, M. J., Goldsmith, J. A., Hsieh, C. L., Johnson, N. V., Wrapp, D., Mclellan, J. S., Barnhart, B. C., Graham, B. S., Mascola, J. R., Hansen, C. L. & Falconer, E. 2020. LY-CoV555, a rapidly isolated potent neutralizing antibody, provides protection in a non-human primate model of SARS-CoV-2 infection. bioRxiv.

Liu, J., Liu, Y., Xia, H., Zou, J., Weaver, S. C., Swanson, K. A., Cai, H., Cutler, M., Cooper, D., Muik, A., Jansen, K. U., Sahin, U., Xie, X., Dormitzer, P. R. & Shi, P. Y. 2021a. BNT162b2-elicited neutralization of B.1.617 and other SARS-CoV-2 variants. Nature.

Liu, L., Wang, P., Nair, M. S., Yu, J., Rapp, M., Wang, Q., Luo, Y., Chan, J. F., Sahi, V., Figueroa, A., Guo, X. V., Cerutti, G., Bimela, J., Gorman, J., Zhou, T., Chen, Z., Yuen, K. Y., Kwong, P. D., Sodroski, J. G., Yin, M. T., Sheng, Z., Huang, Y., Shapiro, L. & Ho, D. D. 2020. Potent neutralizing antibodies against multiple epitopes on SARS-CoV-2 spike. Nature, 584, 450–456.

Liu, Z., Vanblargan, L. A., Bloyet, L. M., Rothlauf, P. W., Chen, R. E., Stumpf, S., Zhao, H., Errico, J. M., Theel, E. S., Liebeskind, M. J., Alford, B., Buchser, W. J., Ellebedy, A. H., Fremont, D. H., Diamond, M. S. & Whelan, S. P. J. 2021b. Identification of SARS-CoV-2 spike mutations that attenuate monoclonal and serum antibody neutralization. Cell Host Microbe, 29, 477–488 e4.

Mccallum, M., De Marco, A., Lempp, F. A., Tortorici, M. A., Pinto, D., Walls, A. C., Beltramello, M., Chen, A., Liu, Z., Zatta, F., Zepeda, S., Di Iulio, J., Bowen, J. E., Montiel-Ruiz, M., Zhou, J., Rosen, L. E., Bianchi, S., Guarino, B., Fregni, C. S., Abdelnabi, R., Foo, S. C., Rothlauf, P. W., Bloyet, L. M., Benigni, F., Cameroni, E., Neyts, J., Riva, A., Snell, G., Telenti, A., Whelan, S. P. J., Virgin, H. W., Corti, D., Pizzuto, M. S. & Veesler, D. 2021. N-terminal domain antigenic mapping reveals a site of vulnerability for SARS-CoV-2. Cell, 184, 2332–2347 e16.

Plante, J. A., Liu, Y., Liu, J., Xia, H., Johnson, B. A., Lokugamage, K. G., Zhang, X., Muruato, A. E., Zou, J., Fontes-Garfias, C. R., Mirchandani, D., Scharton, D., Bilello, J. P., Ku, Z., An, Z., Kalveram, B., Freiberg, A. N., Menachery, V. D., Xie, X., Plante, K. S., Weaver, S. C. & Shi, P. Y. 2021a. Spike mutation D614G alters SARS-CoV-2 fitness. Nature, 592, 116–121.

Plante, J. A., Mitchell, B. M., Plante, K. S., Debbink, K., Weaver, S. C. & Menachery, V. D. 2021b. The variant gambit: COVID-19’s next move. Cell Host Microbe, 29, 508–515.

Polack, F. P., Thomas, S. J., Kitchin, N., Absalon, J., Gurtman, A., Lockhart, S., Perez, J. L., Perez Marc, G., Moreira, E. D., Zerbini, C., Bailey, R., Swanson, K. A., Roychoudhury, S., Koury, K., Li, P., Kalina, W. V., Cooper, D., Frenck, R. W., JR., Hammitt, L. L., Tureci, O., Nell, H., Schaefer, A., Unal, S., Tresnan, D. B., Mather, S., Dormitzer, P. R., Sahin, U., Jansen, K. U., Gruber, W. C. & Group, C. C. T. 2020. Safety and Efficacy of the BNT162b2 mRNA Covid-19 Vaccine. N Engl J Med, 383, 2603–2615.

Riepler, L., Rossler, A., Falch, A., Volland, A., Borena, W., Von Laer, D. & Kimpel, J. 2020. Comparison of Four SARS-CoV-2 Neutralization Assays. Vaccines (Basel), 9.

Schmidt, F., Weisblum, Y., Muecksch, F., Hoffmann, H. H., Michailidis, E., Lorenzi, J. C. C., Mendoza, P., Rutkowska, M., Bednarski, E., Gaebler, C., Agudelo, M., Cho, A., Wang, Z., Gazumyan, A., Cipolla, M., Caskey, M., Robbiani, D. F., Nussenzweig, M. C., Rice, C. M., Hatziioannou, T. & Bieniasz, P. D. 2020. Measuring SARS-CoV-2 neutralizing antibody activity using pseudotyped and chimeric viruses. J Exp Med, 217.

Shi, R., Shan, C., Duan, X., Chen, Z., Liu, P., Song, J., Song, T., Bi, X., Han, C., Wu, L., Gao, G., Hu, X., Zhang, Y., Tong, Z., Huang, W., Liu, W. J., Wu, G., Zhang, B., Wang, L., Qi, J., Feng, H., Wang, F. S., Wang, Q., Gao, G. F., Yuan, Z. & Yan, J. 2020. A human neutralizing antibody targets the receptor-binding site of SARS-CoV-2. Nature, 584, 120–124.

Singh, J., Rahman, S. A., Ehtesham, N. Z., Hira, S. & Hasnain, S. E. 2021. SARS-CoV-2 variants of concern are emerging in India. Nat Med.

Starr, T. N., Greaney, A. J., Dingens, A. S. & Bloom, J. D. 2021. Complete map of SARS-CoV-2 RBD mutations that escape the monoclonal antibody LY-CoV555 and its cocktail with LY-CoV016. Cell Rep Med, 2, 100255.

Suryadevara, N., Shrihari, S., Gilchuk, P., Vanblargan, L. A., Binshtein, E., Zost, S. J., Nargi, R. S., Sutton, R. E., Winkler, E. S., Chen, E. C., Fouch, M. E., Davidson, E., Doranz, B. J., Chen, R. E., Shi, P. Y., Carnahan, R. H., Thackray, L. B., Diamond, M. S. & Crowe, J. E., JR. 2021. Neutralizing and protective human monoclonal antibodies recognizing the N-terminal domain of the SARS-CoV-2 spike protein. Cell, 184, 2316–2331 e15.

Tian, S., Hu, W., Niu, L., Liu, H., Xu, H. & Xiao, S. Y. 2020. Pulmonary Pathology of Early-Phase 2019 Novel Coronavirus (COVID-19) Pneumonia in Two Patients With Lung Cancer. J Thorac Oncol, 15, 700–704.

Wall, E. C., Wu, M., Harvey, R., Kelly, G., Warchal, S., Sawyer, C., Daniels, R., Hobson, P., Hatipoglu, E., Ngai, Y., Hussain, S., Nicod, J., Goldstone, R., Ambrose, K., Hindmarsh, S., Beale, R., Riddell, A., Gamblin, S., Howell, M., Kassiotis, G., Libri, V., Williams, B., Swanton, C., Gandhi, S. & Bauer, D. L. 2021. Neutralising antibody activity against SARS-CoV-2 VOCs B.1.617.2 and B.1.351 by BNT162b2 vaccination. Lancet, 397, 2331–2333.

Wang, R., Chen, J., Gao, K. & Wei, G. W. 2021. Vaccine-escape and fast-growing mutations in the United Kingdom, the United States, Singapore, Spain, India, and other COVID-19-devastated countries. Genomics, 113, 2158–2170.

Xia, S., Zhang, Y., Wang, Y., Wang, H., Yang, Y., Gao, G. F., Tan, W., Wu, G., Xu, M., Lou, Z., Huang, W., Xu, W., Huang, B., Wang, H., Wang, W., Zhang, W., Li, N., Xie, Z., Ding, L., You, W., Zhao, Y., Yang, X., Liu, Y., Wang, Q., Huang, L., Yang, Y., Xu, G., Luo, B., Wang, W., Liu, P., Guo, W. & Yang, X. 2021. Safety and immunogenicity of an inactivated SARS-CoV-2 vaccine, BBIBP-CorV: a randomised, double-blind, placebo-controlled, phase 1/2 trial. Lancet Infect Dis, 21, 39–51.

Xu, Z., Shi, L., Wang, Y., Zhang, J., Huang, L., Zhang, C., Liu, S., Zhao, P., Liu, H., Zhu, L., Tai, Y., Bai, C., Gao, T., Song, J., Xia, P., Dong, J., Zhao, J. & Wang, F. S. 2020. Pathological findings of COVID-19 associated with acute respiratory distress syndrome. Lancet Respir Med, 8, 420–422.

Zhou, B., Thao, T. T. N., Hoffmann, D., Taddeo, A., Ebert, N., Labroussaa, F., Pohlmann, A., King, J., Steiner, S., Kelly, J. N., Portmann, J., Halwe, N. J., Ulrich, L., Trueb, B. S., Fan, X., Hoffmann, B., Wang, L., Thomann, L., Lin, X., Stalder, H., Pozzi, B., De Brot, S., Jiang, N., Cui, D., Hossain, J., Wilson, M. M., Keller, M. W., Stark, T. J., Barnes, J. R., Dijkman, R., Jores, J., Benarafa, C., Wentworth, D. E., Thiel, V. & Beer, M. 2021. SARS-CoV-2 spike D614G change enhances replication and transmission. Nature, 592, 122–127.

